# Age-related differences in the influence of task-irrelevant information on the neural bases of phonological and semantic processes

**DOI:** 10.1101/136606

**Authors:** Michele T. Diaz, Micah A. Johnson, Deborah M. Burke, Trong-Kha Truong, David J. Madden

## Abstract

Although aging is often associated with cognitive decline, there is considerable variability among individuals and across domains of cognition. Within language, several indicators of semantic processing show stability throughout the lifespan. However, older adults have increased difficulty with phonological aspects of language, especially in language production. While these behavioral patterns are established, the neurobiology associated with these behaviors are less clear. Previously we have shown that, older adults were slower and less accurate in phonological compared to semantic decisions, and that older adults didn’t exhibit brain-behavior relationships. In the present study, we examined phonological and semantic processes in the presence of task-irrelevant information. Older and younger adults made phonological and semantic decisions about pictures in the presence of either phonologically-related or semantically-related words, which were unrelated to the task. Behavioral results indicated that overall older adults had slower reaction times and lower accuracy compared to younger adults, and that all adults were less efficient when making phonological compared to semantic decisions. Patterns of brain activation for the semantic condition showed that all adults engaged typical left-hemisphere language regions, and that this activation was positively correlated with efficiency. In contrast, the phonological condition elicited activation in bilateral precuneus and cingulate, but only younger adults showed a significant relationship between activation and efficiency. Our results suggest that the relationship between behavior and neural activation when processing phonological information declines with age, but that the core semantic system continues to be engaged throughout the lifespan, even in the presence of task-irrelevant information.

## INTRODUCTION

Although aging is often characterized by cognitive decline, there is considerable variability across individuals and domains. For example, in the area of language production, one of the most commonly reported challenges for older adults is word retrieval (for a review see Burke & Shafto, 2008). These retrieval deficits are hypothesized to arise, at least in part, from impaired phonological retrieval. Consistent with this hypothesis, older adults report an increased incidence of tip of the tongue experiences, where one knows a word but is unable to produce the phonology (Brown & McNeill, 1966; Burke, Mackay, Worthley, & Wade, 1991). In contrast, older adults tend to show comparable performance, relative to younger adults, in most forms of semantic retrieval. Older and younger adults have similar levels of semantic priming (Bowles, Williams, & Poon, 1983; Burke, White, & Diaz, 1987; Madden, Pierce, & Allen, 1993); older adults exhibit stable or increased vocabulary scores across the lifespan (Kemper & Sumner, 2001; Singer, Verhaeghen, Ghisletta, Lindenberger, & Baltes, 2003; Verhaeghen, 2003); and older adults produce more lexically diverse utterances (Kemper & Sumner, 2001). Each of these observations points to a larger and more elaborate semantic system for older adults.

Core phonological and semantic processes, however, interact with other aspects of cognition, such as attention and working memory, which tend to show age-related decline. Hasher and Zacks (1988) proposed the inhibition deficit hypothesis to explain why older adults are slowed more than younger adults when text is interspersed with to-be-ignored text, particularly when the distracting text is related to the other presented material (Connelly, Hasher, & Zacks, 1991), or presented in an unpredictable location (Carlson, Hasher, Zacks, & Connelly, 1995). The potential interaction between hypothesized age-related inhibitory declines and language production, and the neural bases for these effects, are unknown.

Previous research from our lab comparing phonological and semantic decisions has shown that older adults were less accurate and less efficient in making phonological judgments about pictures, whereas performance did not differ for semantic judgments (Diaz, Johnson, Burke, & Madden, 2014). Analyses of the fMRI activation related to these decisions showed that while the semantic task engaged typical left hemisphere language regions, the phonological task engaged bilateral cingulate and precuneus, suggesting a greater reliance on task-control regions during the phonological task. Consistent with previous findings, older adults elicited greater activation than younger adults throughout the brain. However, these increases in activation were not related to behavioral performance for older adults, suggesting a disconnection between the neural response and behavioral performance.

Several other studies have also examined age-related differences in the neural bases of language production. These studies contribute to our understanding of whether age-related differences in activation follow compensatory (e.g., HAROLD, Cabeza, Anderson, Locantore, & McIntosh, 2002) or dedifferentiated (e.g., Li, Lindenberger, & Sikstrom, 2001) patterns in language production. The earliest reports, examining picture naming, largely supported a compensatory pattern, showing that older adults demonstrated similar accuracy compared to younger adults while eliciting increased activation in right inferior frontal gyrus and bilateral cingulate (Wierenga et al., 2008). Moreover, older adults’ increased activation in right inferior frontal gyrus was positively correlated with accuracy in Wierenga et al., 2008. Similarly, Shafto and colleagues reported that older adults’ increased activation in the insula during word retrieval was associated with fewer tip of the tongue states (i.e., increases in activation were associated with fewer retrieval failures Shafto, Stamatakis, Tam, & Tyler, 2010). Effects consistent with compensation have also been reported for a task requiring rhyme judgments about words (Geva et al., 2012), in which activation in right inferior frontal gyrus (IFG) increased with age and older and young adults responded with similar speed. However, older adults who had the highest error rates elicited the greatest right IFG activation, suggesting that increased activation may reflect compensatory effort, but not necessarily successful performance. Interestingly, a split-half analysis of higher and lower performing older adults in the Wierenga et al., 2008 picture naming study showed opposite patterns: High performers had a positive correlation between accuracy and rIFG fMRI activation, while lower performers had a negative relation between these variables (Wierenga et al., 2008). These findings suggest that there are complex relations between age-related differences in activation and behavioral performance, and that more than one neurocognitive profile may exist.

Here we used fMRI and behavioral measures to investigate the influence of additional information on phonological, semantic, and perceptual judgments in younger and older adults. Participants made phonological, semantic, and perceptual judgments about two photographs, presented with phonologically-related, semantically-related, or non-word distractor letter strings. Photos were used for the target stimuli so as not to provide explicit orthographic or phonological input about the item to be recalled, and across phonological and semantic tasks identical photographs were included to control for perceptual features. In contrast, we included words and non-words as the task-irrelevant stimuli to provide orthographically and phonologically salient task-irrelevant information.

On the basis of these previous findings, we predicted that if older adults are less efficient than younger adults in the inhibition of irrelevant information, then we would expect large age-related differences for both phonological and semantic conditions. However, if younger and older adults process this additional information in a similar manner, this should result in a reduction of age-related differences. Along similar lines, if additional information is detrimental to older adults’ performance, then we would expect age-related differences in the neural response, particularly in regions associated with executive function and task control. However, if older adults process this information similarly to younger adults, then we may expect to see similar patterns of activation across age groups. Finally, we examined age differences in the relation between neural mechanisms and behavior, and how these relationships may differ across conditions. Specifically, we examined the results for patterns of neural compensation or inefficiency in older adults. Increases in activation for older adults that correspond to maintained or improved behavior would be consistent with a compensation account. Whereas weaker or no relation between fMRI activation and behavioral performance in older adults would be consistent with neural inefficiency.

## METHODS

### Participants

All participants were community-dwelling, right-handed, native English speakers. There were 20 younger adults (*M* age 25.0; age range 19-35; 8 male) and 20 older adults (*M* age 67.3; age range 59-76; 8 male). One younger participant was removed for a high number of omitted responses during the task, and one older participant was removed due to experiencing anxiety in the scanner, leaving 19 participants in each group. All participants had normal or corrected to normal vision by self-report. None was color blind, reported a history of neurological or psychological disorders, or any major medical conditions (e.g., diabetes, heart disease, Christensen, Moye, Armson, & Kern, 1992). All participants completed neuropsychological testing to assess basic cognitive skills such as speed, memory, executive function, and language. Across groups, participants did not differ in years of education, MMSE scores, measures of anxiety and depression, vocabulary, verbal fluency, digit symbol forward, or immediate recall. Demographic characteristics are presented in Table 1. Each participant provided informed consent and was paid for his or her participation. All experimental procedures were approved by the Duke University School of Medicine Institutional Review Board.

**Table 1:** Participant Demographics

|  | Younger | Older |
| --- | --- | --- |
| N | 19 | 19 |
| Age*** | 25.0 (4.90) | 67.47 (5.36) |
| Education | 16.37 (2.14) | 17.16 (2.01) |
| MMSE | 29.16 (1.01) | 29.26 (0.81) |
| HADS | 0.05 (0.08) | 0.07 (0.08) |
| Vocabulary (WAIS II) | 57.63 (5.74) | 57.37 (4.94) |
| Verbal Fluency (total) | 72.32 (18.45) | 66.37 (16.57) |
| Digit Symbol (DS) RT*** | 1271.48 (239.16) | 1861.49 (350.84) |
| DS Forward | 12.63 (2.71) | 11.42 (2.39) |
| DS Backward** | 9.63 (2.67) | 7.68 (1.83) |
| Stroop RT*** | 492.42 (88.54) | 647.40 (167.56) |
| Speed RT*** | 294.87 (40.07) | 356.25 (64.02) |
| Immediate Recall | 11.68 (3.04) | 10.79 (2.74) |
| Delayed Recall* | 10.89 (3.65) | 9.00 (3.32) |

Values provided are means and (SD). \* denotes $p < .05$ , \*\* denotes $p < .01$ , \*\*\* denotes $p < .0001$

### Stimuli

On each trial, participants made one of three types of judgments: phonological, semantic, or perceptual (a control condition). Figure 1 provides an overview of the task design. Stimuli consisted of a cue (phonological, semantic, or perceptual) followed by a pair of photographs and a centrally-presented distractor word that was unrelated to the task. Cues were a word or short statement followed by a question mark. Participants were asked to decide whether or not both photographs matched the cue. For half of the trials, both pictures matched the cue (i.e., match trials) and for the other half only one picture matched the cue (i.e., non-match trials). Phonological cues presented a question about the first letter of the object names (e.g., Starts with B?, Starts with P?)^1^. Semantic cues presented a question about a functional or perceptual attribute of the objects (e.g., Smooth?, Flies?, Edible?). Perceptual cues always presented a question about whether the two items were identical (e.g., Same?). Photographs were high-resolution images of everyday objects against a white background. On phonological trials, the distractor word rhymed with one of the pictures. On semantic trials, the distractor word was semantically-related to one of the objects depicted in the photographs. Perceptual distractor stimuli were random consonant strings. In this way, the distractor stimuli were always related to one of the objects (phonologically, semantically, or perceptually), but did not convey information that facilitated the decision, and the relationship of the distractor stimuli to the photo pairs was always the same regardless of whether the photo pairs matched the cue or not.

Across conditions, the names of objects did not statistically differ in length (number of letters; phonological match = 5.50; phonological non-match = 5.56; semantic match = 5.51; semantic non-match = 5.59), number of syllables (phonological match = 1.58; phonological non-match = 1.65; semantic match = 1.62; semantic non-match = 1.65), number of phonemes (phonological match = 4.52; phonological non-match = 5.02; semantic match = 4.57; semantic non-match = 4.64), or word frequency (SUBTL; phonological match = 37.32; phonological non-match = 32.20; semantic match = 31.40; semantic non-match = 35.17). Although not a phonological or semantic feature per se, frequency was controlled because of its well-established influence on naming (e.g., Oldfield & Wingfield, 1965). Ratings of frequency were obtained from the English Lexicon Project (Balota et al., 2007). Distractor words did not statistically differ across conditions in either length or word frequency.

During functional magnetic resonance imaging (fMRI) scanning, the entire experiment comprised 240 trials, with 80 trials (40 match, 40 non-match) in each of the three judgment conditions. Trial types were evenly distributed across the 8 runs in a random ordering. Individual objects occurred once for each participant but were counterbalanced across the phonological and semantic conditions across participants, with two separate trial lists. The perceptual control condition served to control for basic visual and motor processes. We created images for the perceptual judgments in Matlab (Mathworks, Natick, MA USA) by applying a Fourier transform to a random subset of the object photographs, permuting the phase spectrum, and then computing an inverse transform. We then adjusted the luminance of the perceptual photographs to match the original photograph. This procedure yielded displays that contained the basic perceptual features of the original photographs, but the features did not combine to form a recognizable object. Examples of the original and perceptual images can be found in Supplemental Figure 1. All stimuli were selected based on two separate behavioral pretests, as we have previously reported (Diaz et al., 2014).

**Figure 1.**
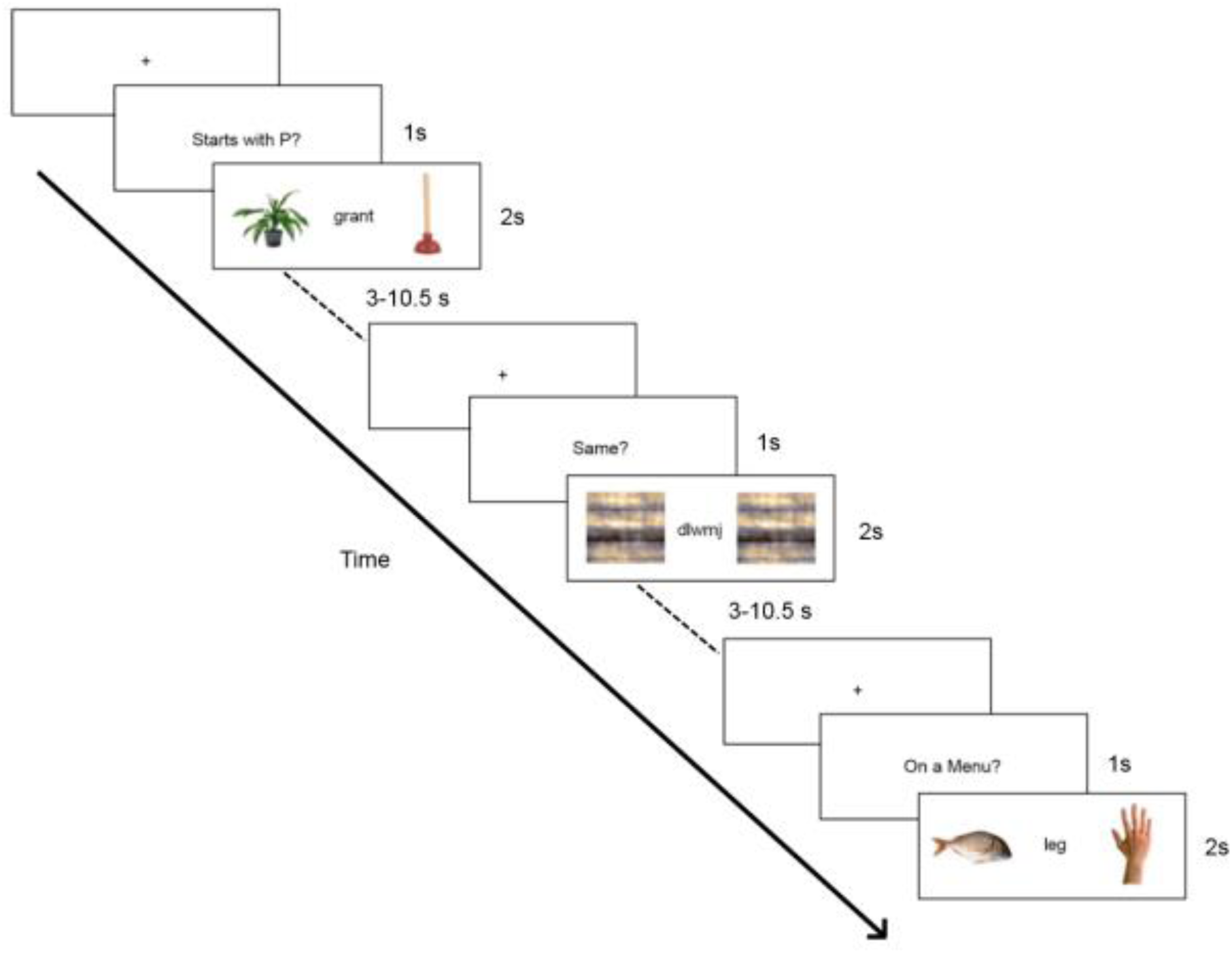
Task Design. An overview of the task design is provided. Here we highlight typical trials (semantic, perceptual, and phonological). Cues were presented for 1 s followed by pictures and distractor words that were presented for 2 s. The inter-stimulus interval was jittered between 3-10.5 s with a mean interval of 4.8 s.

### Experimental Task

Each trial consisted of a cue (duration = 1 s) followed by two photographs of objects presented side by side (duration = 2 s). The two objects either both matched the cue or one item was incongruent with the cue. Simultaneously presented with the photographs was a written word that was phonologically or semantically related to one of the photographs (i.e., rhymed or from the same semantic category). Participants were asked to decide if both photographs matched the cue and made their match/non-match response manually with the index and middle finger of the right hand. They were instructed to respond at the onset of the object photographs as quickly as possible while maintaining accuracy. The three conditions: phonological, semantic, or perceptual, combined orthogonally with match/non-match to yield six different trial types (Figure 1).

All six trial types were presented in a randomized order within each fMRI run, with a variable inter-trial-interval (ITI; interval = 3 - 10.5 s, *M* = 4.8 s). Trial order across conditions and inter-trial-intervals were randomized and optimized using the Optseq2 program (Dale, 1999). Trials were distributed across eight runs of approximately four min, each of which began and ended with the presentation of a fixation cross for 6 s and 15 s, respectively. A fixation cross was also presented during the variable interval between each trial. All stimuli were presented via a projector using an in-house experimental control program (Voyvodic, 1999). Responses were recorded with a hand-held fiber optic response box (Current Designs, Philadelphia, PA, USA).

### Acquisition of MRI Data

MRI scanning was completed on a 3.0 Tesla GE MR 750 whole-body 60 cm bore human scanner equipped with 50 mT/m gradients and a 200 T/m/s slew rate. An eight-channel head coil was used for Radio Frequency reception (General Electric, Milwaukee, WI USA). Sagittal T-1 weighted localizer images were acquired and used to define a volume for data collection and high order shimming. The anterior and posterior commissures were identified for slice selection and shimming. A semi-automated high-order shimming program was used to ensure global field homogeneity. High-resolution structural images were acquired using a 3D fSPGR pulse sequence (TR = 8.14 ms; TE = 3.22 ms; TI = 450 ms; FOV = 24 cm^2^; flip angle =12°; voxel size = 0.9375 × 0.9375 × 1mm; matrix = 256 × 256; 162 contiguous slices; averages = 1; Phase Encoding: RL; bandwidth: 62.5 kHz). Functional images sensitive to blood oxygen level-dependent (BOLD) contrast were acquired using an inverse spiral pulse sequence with SENSE acceleration (TR = 2.0s; TE = 30 ms; FOV = 25.6 cm^2^; flip angle = 60°; SENSE factor = 2; voxel size = 3.75 × 3.75 × 4 mm; matrix = 64 × 64; 38 contiguous oblique axial slices, parallel to the AC-PC line, interleaved acquisition; bandwidth: 250 kHz). Four initial RF excitations were performed to achieve steady state equilibrium and were subsequently discarded.

### fMRI Data Analysis

Data were analyzed for quality via a tool that quantifies several metrics including Signal-to-Noise (SNR), Signal-Fluctuation-to-Noise (SFNR), motion, and voxel-wise standard deviation measurements (Friedman & Glover, 2006; Glover et al., 2012). Additionally, all data were visually inspected for artifacts and blurring. The average movement in the X, Y, or Z directions was .23 mm (range .04 – 1.25 mm). Thus, none of the included participants moved more than 1/2 voxel in the X, Y, or Z dimensions. We used FSL version 5.0.1 and FEAT version 6.00 for preprocessing and for all analyses of functional activations (Smith et al., 2004; Woolrich et al., 2009). Pre-whitening or voxel-wise temporal autocorrelation was estimated and corrected using FILM, FMRIB’s Improved Linear Model (Woolrich, Ripley, Brady, & Smith, 2001). The skull and other coverings were stripped from the structural brain images using the FSL brain extraction tool (Smith, 2002). Functional image data were corrected for slice timing using sinc interpolation to shift each slice in time to the middle of the TR period. Functional images were motion-corrected using FSL’s MC-FLIRT (FMRIB’s Linear Image Registration Tool) using 6 rigid-body transformations (Jenkinson, Bannister, Brady, & Smith, 2002). These estimates of motion were included as nuisance covariates in the overall FSL model. Functional data were also high-pass filtered (cut off = 50s), and spatially smoothed using a Gaussian kernel (FWHM = 8 mm). Functional images of each participant were co-registered to structural images in native space, and structural images were normalized to Montreal Neurological Institute (MNI) standard space using FSL’s MNI Avg152 T1 2 × 2 × 2 mm standard brain. The same transformation matrices used for structural-to-standard transformations were then used for functional-to-standard space transformations of co-registered functional images. Co-registration and normalization steps were completed using a combination of affine and non-linear registrations (Greve & Fischl, 2009; Jenkinson et al., 2002; Jenkinson & Smith, 2001).

We used a double γ function to model the hemodynamic response for each trial. Correct trials were modeled by condition, and errors (incorrect responses and omitted responses) were modeled as a separate regressor. Reaction time (RT) outliers were defined as responses < 250 ms or ± 3 SD from that individual’s overall mean. These accounted for approximately 9% of the trials (7.2% incorrect, 0.47% no response, 0.95% outliers).

We combined the analyses from each experimental run and performed an analysis across runs for each participant individually. We then combined these analyses across participants into a group level analyses using the FMRIB Local Analysis of Mixed Effects (FLAME 1 & 2) to identify voxels that were activated by each condition (Beckmann, Jenkinson, & Smith, 2003; Woolrich, Behrens, Beckman, Jenkinson, & Smith, 2004). Our primary analytic goals were to identify regions that were responsive to our experimental manipulation (phonological and semantic processing), to identify regions where older and younger adults differed, and to detect regions in which the activation differed as a function of both age group and task condition concurrently (i.e., interaction). To accomplish this, we performed a two-way analysis of variance (ANOVA) within FSL with Condition (Phonological, Semantic), Age Group (Younger, Older), and the interaction of Condition × Age Group as independent variables. Because perceptual trials were qualitatively different from the language conditions they were omitted from this ANOVA. Within FSL, we also made comparisons between conditions to identify differences in functional activation between levels of each variable (e.g., Phonological > Semantic, Semantic > Phonological). All significant activations were determined using a two-step process in which voxels, significant at *p* < .01, were identified. Clusters of identified voxels were then corrected for multiple comparisons according to Gaussian random fields (GRF) theory (*p* < .05, corrected) in which each cluster’s estimated significance level was compared with the cluster probability threshold, and then only clusters whose estimated significance exceeded the threshold were included in the results (Hayasaka & Nichols, 2003). Results from comparisons between conditions (e.g., phonological > semantic) were masked by results from more basic analyses (e.g., significant activation to phonological trials alone) in a conjunction analysis to focus the activation contrasts on positive HDR responses. All analyses involved a whole-brain approach with single comparisons, and thus the comparisons should not be statistically biased (Kriegeskorte, Simmons, Bellgowan, & Baker, 2009). We determined anatomical gyri corresponding to the peaks of activation through reference to anatomical atlases (Desikan et al., 2006). All reported coordinates are in MNI space, and results are overlaid on the MNI template brain.

## RESULTS

### Behavioral Results

Behavioral data are presented in Figure 2. For all of our analyses of behavioral results we performed univariate analyses of variance (ANOVAs) with Age Group as a between-subjects variable and Condition (phonological, semantic) and Match type (match, non-match) as within-subjects variables. Because the perceptual task was qualitatively different from the other two conditions, we did not include it in our analyses of behavioral results.

**Figure 2.**
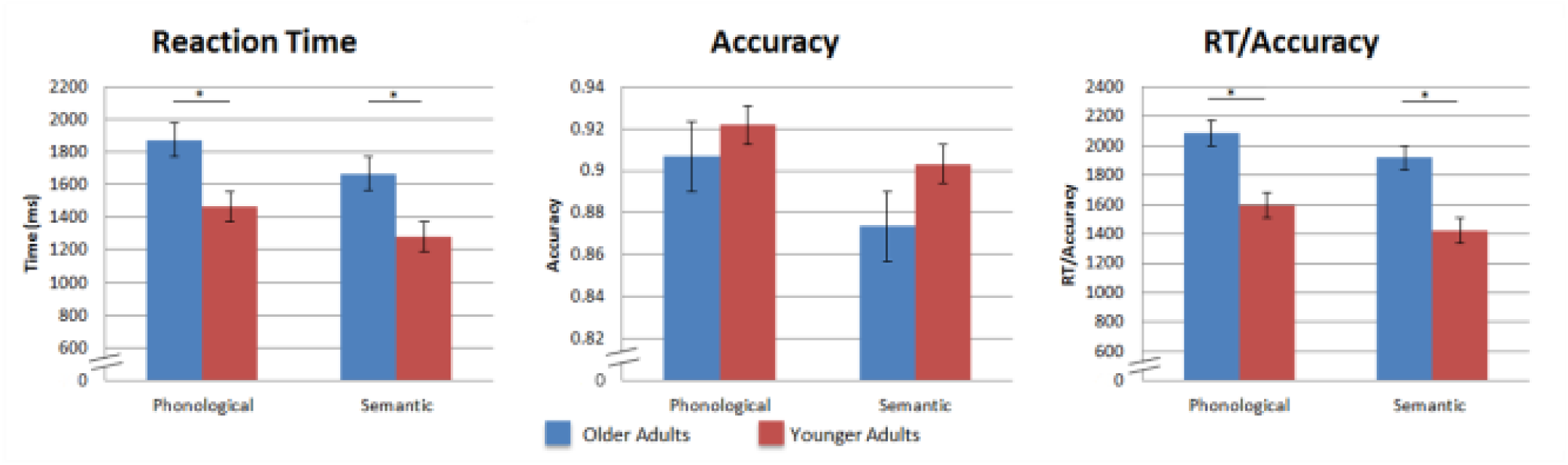
Behavioral Results. Shown are group means for reaction time (RT), accuracy, and RT/Accuracy with standard error bars. There was a main effect of Condition in all three behavioral measures with participants responding more slowly and more accurately to the phonological trials and responding faster but less accurately to the semantic trials. There was also a main effect of Age Group in the RT and RT/accuracy measures with older adults responding more slowly overall. There were no significant interactions between Age Group and Condition.

#### Reaction time

Analysis of the mean untransformed RT data for correct responses indicated that there was a significant main effect of Age Group in which older adults were slower than younger adults, *F*(1, 36) = 12.14, *p* < .01. There was an additional main effect of Condition in which participants responded more quickly in the semantic condition than in the phonological condition, *F*(1,36) = 80.24, *p* < .0001. We also observed a significant main effect of Match, *F*(1,36) = 27.44, *p* < .0001, and a significant Condition × Match interaction, *F*(1,36) = 45.73, *p* < .0001. Overall, match responses were faster than non-match responses, and this difference was significant in the phonological condition, *F*(1,36) = 46.04, *p* < .0001, but not in the semantic condition.

#### Accuracy

Analysis of accuracy indicated that there was not a significant main effect of Age Group, and there were no significant interactions with Age Group. There was a significant main effect of Condition in which participants were more accurate in the phonological condition than the semantic condition, *F*(1,36) = 9.88, *p* < .01. We also observed a significant Condition × Match interaction, *F*(1,36) = 49.15, *p* < .0001. Overall, there were no differences in accuracy for match and non-match responses. However, in the phonological condition participants were significantly more accurate on match trials, *F*(1,36) = 4.86, *p* < .05, and the opposite was observed in the semantic condition where participants were significantly more accurate on non-match trials, *F*(1,36) = 26.02, *p* < .0001.

#### RT/Accuracy

Because there appeared to be speed accuracy trade-offs that differed as a function of condition (i.e., slower but more accurate performance in the phonological condition relative to the semantic condition) we also conducted an analysis of RT divided by accuracy. Essentially, this analytic approach adjusts RT to incorporate accuracy and can serve as a measure of overall processing efficiency, with higher values representing less efficient performance, similar to RT (e.g., Horowitz & Wolfe, 2003; Townsend & Ashby, 1983). This analysis indicated that efficiency was worse for older adults than for younger adults, *F*(1,36) = 15.22, *p* < .0005, and was worse in the phonological condition than the semantic condition *F*(1,36) = 30.5, *p* < .0001. We also observed a significant main effect of Match, *F*(1,36) = 310.86, *p* < .005, and a significant Condition × Match interaction, *F*(1,36) = 88.25, *p* < .0001. In the phonological condition, performance was less efficient on non-match relative to all other trials, and there were no significant differences in efficiency across the other three conditions. Finally, there was a significant Age Group × Condition × Match interaction, *F*(1,36) = 6.03, *p* < .05. Both groups showed similar patterns in which performance on phonological non-match trials was less efficient than all other conditions, and this difference was larger for older adults than for younger adults. Thus, with the exception of the Age Group × Condition X Match interaction, the RT/accuracy results largely mirrored our reaction time results.

**Figure 3.**
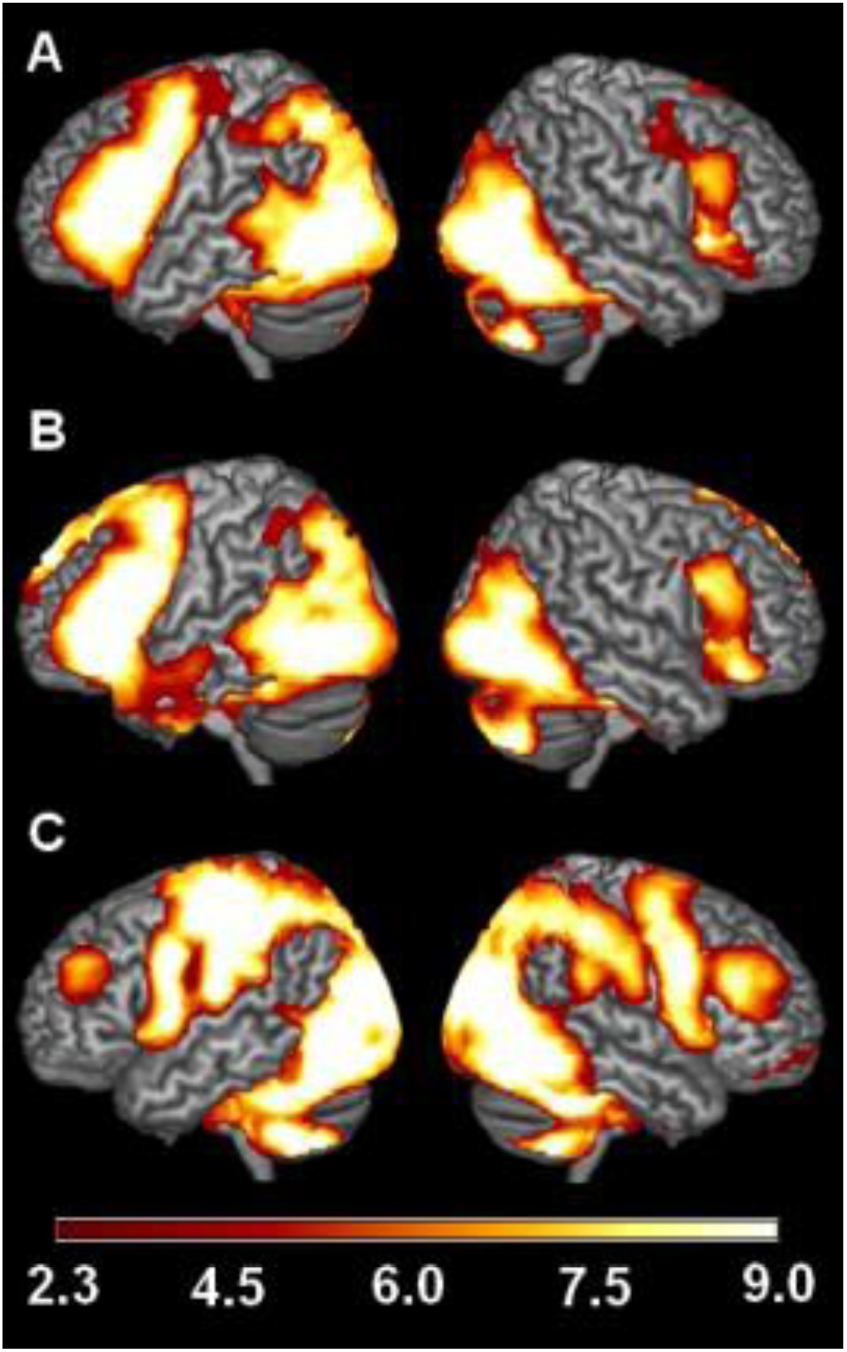
fMRI Activations to Individual Conditions An overview of the regions in which there was significant activation comparing a) phonological trials > perceptual control trials, b) semantic trials > perceptual control trials, and c) the perceptual control trials > null events. Colored regions represent areas where significant differences were found, p<.01 corrected.

### Comparisons with our previous experiment

To further assess the influence of the distractors on behavioral performance, we conducted a comparison between the behavioral results from the present study and those from our earlier experiment (Diaz et al., 2014). There were no significant main effects of study for RT, accuracy, or RT/ACC. However, for accuracy there was a marginally significant interaction of study × group x condition, with older adults from the present experiment performing more accurately on phonological trials compared with older adults from the first experiment which did not have distractors, *F*(1,126) = 3.48, *p* = .065. There were no differences in accuracy for the semantic condition between the two studies for older or younger adults.

### fMRI Results

First, we report the activation to individual conditions compared with our perceptual control condition and activation to our perceptual control condition compared to baseline (Figure 3). Across participants, trials of primary interest (phonological and semantic) elicited greater activation than our perceptual control condition bilaterally in traditional language regions including inferior frontal gyrus, dorsal medial prefrontal cortex, temporal cortex, and parietal cortex. Consistent with previous studies of language, activation was more extensive in the left hemisphere, and for the semantic condition activation was more extensive in anterior temporal cortex and dorsal medial prefrontal cortex. Relative to the implicit baseline, the perceptual control condition elicited activation in motor cortex and the cerebellum, which is consistent with our-right handed motor response, and also in occipital cortex, which is consistent with the visual nature of the stimulus presentation.

**Figure 4.**
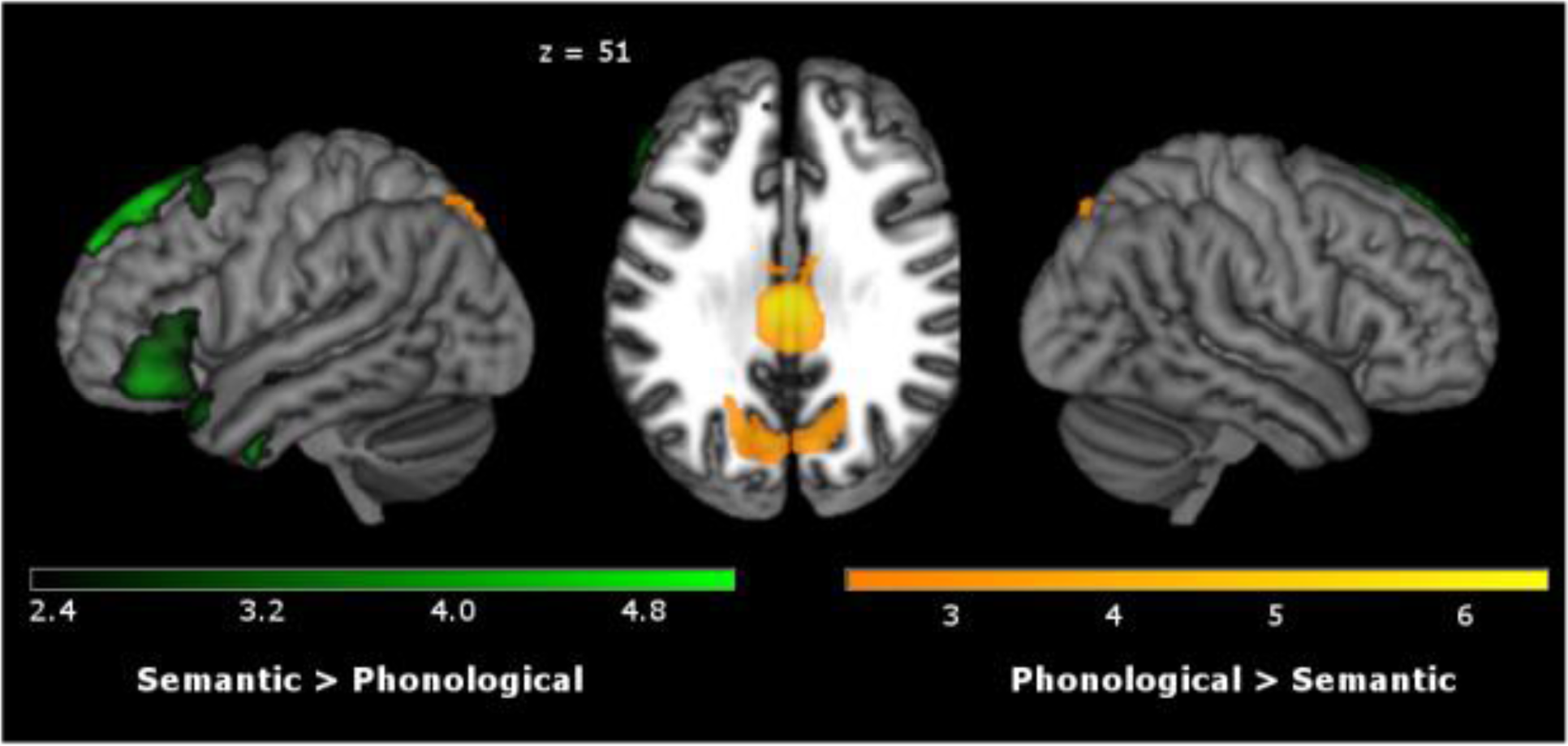
Main Effect of Condition An overview of the regions that comprise the main effect of condition are shown. Regions in which the semantic condition elicited greater activation than the phonological condition are shown in green and regions in which the phonological condition elicited greater activation than the semantic condition are shown in orange. Colored regions represent areas where significant differences were found, p<.01 corrected.

**Table 2:**
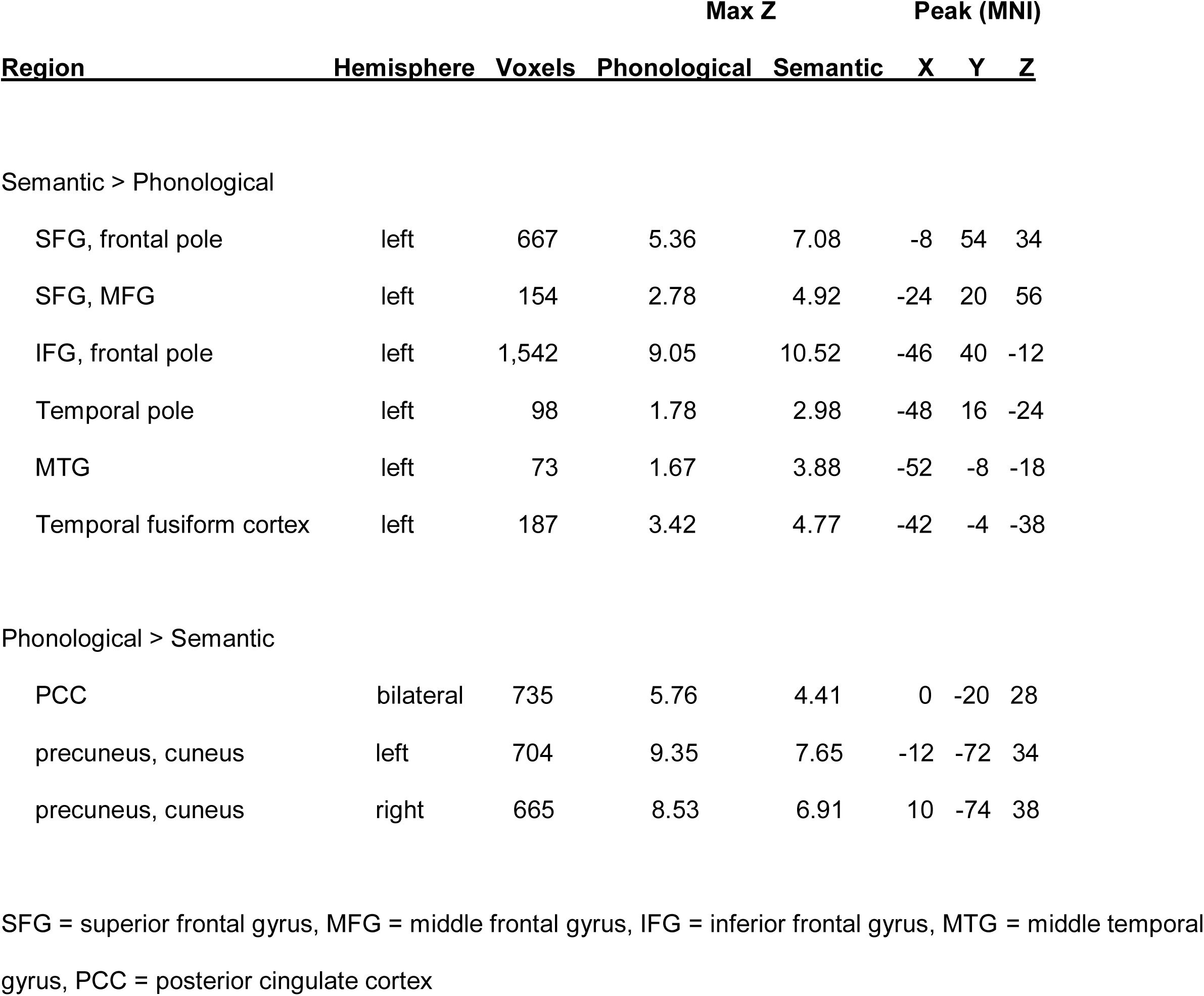
Condition Effects

### Comparisons Between Phonological and Semantic Conditions

Collapsing across age groups, the ANOVA yielded a significant main effect of Condition. The phonological condition elicited greater activation than the semantic condition in bilateral precuneus and bilateral posterior cingulate cortex. Semantic trials elicited greater activation than phonological trials in left hemisphere regions including dorsal medial prefrontal cortex (DMPFC), inferior frontal gyrus (IFG), anterior middle temporal gyrus (MTG), and fusiform gyrus (Figure 4). These results are summarized in Table 2.

**Figure 5.**
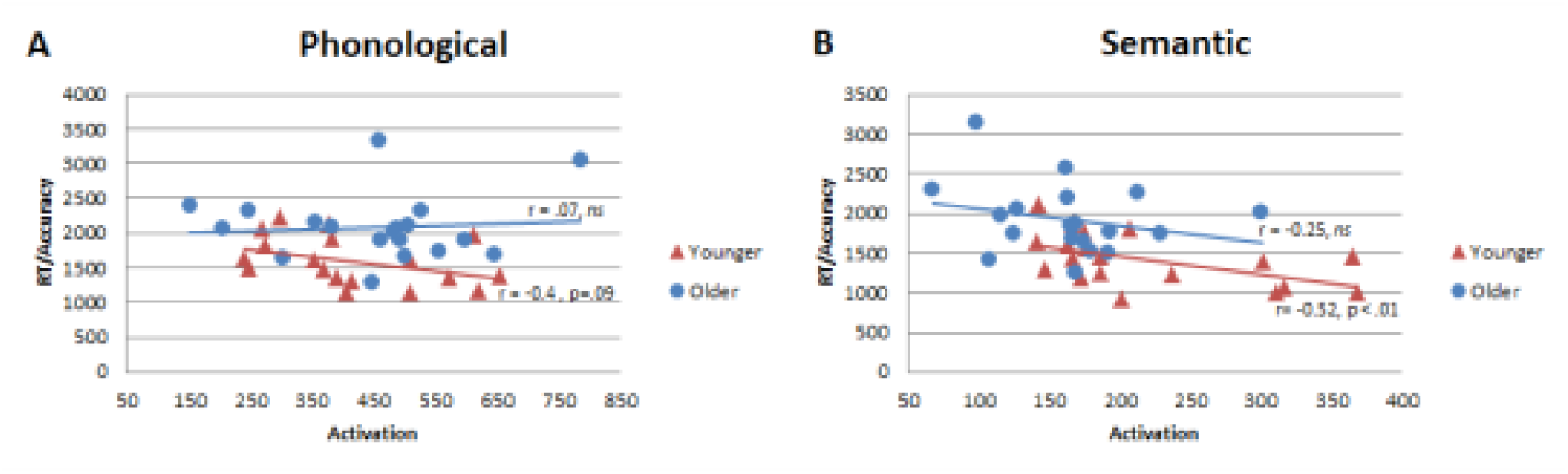
Scatterplot of the Main Effect of Condition Scatterplots of the relationship between our behavioral measure of RT/accuracy and parameter estimates of the fMRI activation from the main effect of Condition are shown. a) For the phonological condition a significant interaction was demonstrated by a marginally significant negative correlation for the younger adults. B) For the semantic condition, there was a significant negative correlation between RT/accuracy and fMRI activation for the overall group.

**Table 3:** Age Effects

| Region | Hemisphere | Voxels | Max Z |  | Peak (MNI) |  |  |
| --- | --- | --- | --- | --- | --- | --- | --- |
|  |  |  | Younger | Older | X | Y | Z |
| Older > Younger |  |  |  |  |  |  |  |
| IFG | left | 1,243 | 13.45 | 13.51 | -42 | 40 | 0 |
| precentral gyrus | left | 48 | 3.30 | 5.50 | -14 | -16 | 64 |
| precentral gyrus | right | 3,622 | 8.18 | 12.28 | 40 | -12 | 64 |
| pre/post -central gyri | left | 986 | 8.39 | 11.98 | -30 | -24 | 46 |
| postcentral gyrus | left | 253 | 6.07 | 8.46 | -46 | -40 | 64 |
| postcentral gyrus | right | 26 | 1.06 | 3.22 | 52 | -18 | 18 |
| supramarginal gyrus | right | 785 | 4.17 | 6.57 | 54 | -24 | 44 |
| supramarginal gyrus | left | 103 | 9.18 | 11.34 | -24 | -52 | 34 |
| anterior cingulate cortex | left | 525 | 5.97 | 10.15 | -2 | -8 | 44 |
| ITG | right | 173 | 11.34 | 11.59 | 48 | -46 | -8 |
| temporal fusiform | left | 2,055 | 16.94 | 17.22 | -36 | -48 | -12 |
| temporal fusiform | right | 681 | 16.45 | 17.21 | 38 | -30 | -24 |
| retrosplenial cortex | bilateral | 843 | 5.52 | 7.99 | 0 | -40 | 14 |
| precuneus, SPL | left | 2,081 | 7.47 | 11.22 | -4 | -54 | 62 |
| precuneus | right | 1,500 | 6.53 | 9.74 | 8 | -60 | 60 |
| SPL | right | 165 | 4.25 | 7.00 | 32 | -44 | 68 |
| occipital cortex | left | 2,149 | 15.89 | 16.73 | -14 | -90 | 2 |
| occipital cortex | right | 1,408 | 16.54 | 17.35 | 28 | -76 | -6 |
| occipital pole | left | 622 | 11.71 | 15.07 | -24 | -96 | 22 |
| occipital pole | right | 592 | 9.38 | 14.21 | 18 | -90 | 38 |
| cerebellum | right | 275 | 15.57 | 16.40 | 34 | -52 | -28 |
| cerebellum | right | 460 | 10.38 | 10.47 | 16 | -74 | -28 |
| thalamus | left | 186 | 2.83 | 4.28 | -4 | 4 | -6 |
| thalamus | right | 223 | 3.83 | 5.44 | 6 | 2 | -6 |

Table 3: Age Effects (continued)
| Region | Hemisphere | Voxels | Max Z |  | Peak (MNI) |  |  |
| --- | --- | --- | --- | --- | --- | --- | --- |
|  |  |  | Younger | Older | X | Y | Z |
| Older > Younger |  |  |  |  |  |  |  |
| brainstem | bilateral | 1,652 | 5.30 | 7.01 | 2 | -30 | -20 |
| Younger > Older |  |  |  |  |  |  |  |
| occipital cortex | left | 380 | 14.73 | 14.48 | -32 | -78 | 20 |

IFG = inferior frontal gyrus, ITG = inferior temporal gyrus, SPL = superior parietal lobule

### Comparisons Between Older and Younger Adults

Collapsing across conditions, the ANOVA yielded a significant main effect of Age Group. Younger adults elicited greater activation than older adults only in occipital cortex. In contrast, older adults elicited greater activation than younger adults in many regions throughout the brain including left IFG, bilateral pre-and post-central gyri, bilateral supramarginal gyri, left anterior cingulate cortex, bilateral precuneus, and bilateral inferior temporal and occipital cortices. The interaction of Condition and Age Group was not significant in the whole brain analysis.

### fMRI Activation-Behavior Relations

To investigate the relationships between activation and behavior we conducted linear regressions in which Age Group, the parameter estimates of the local maxima of fMRI activation, and the interaction of Age Group and fMRI activation were independent variables (predictors), and efficiency (i.e., RT/accuracy) was the outcome variable. Similar analyses conducted with RT yielded similar results and are described in the supplementary materials.

For the phonological condition, the overall model was significant, F(3,36) = 5.18, *p* < .005, R^2^ = 0.31, although no individual predictor was significant. A stepwise regression with the same variables indicated that the interaction term was the strongest predictor of the efficiency measure, F(3,36) = 15.97, *p* < .0005, R^2^ = 0.31, indicating that the relation between activation and efficiency varied across the age groups. The correlation between activation and RT/accuracy was marginally, negatively significant for younger adults *r* = -0.40, *p =* .09 (i.e., greater activation was associated with more efficient phonological decisions, and lower RT/accuracy scores), whereas the correlation was near zero for older adults, *r* = 0.07, n.s.

For the semantic condition, the overall model was significant, F(1,36) = 7.42, *p* < .001, R^2^ = 0.40, and fMRI activation to the semantic condition was also a significant predictor of the efficiency of semantic decisions, F(1,36) = 2.13, *p* < .05. The correlation between RT/accuracy and fMRI activation was negative, *r* = -0.50, *p* < .005. Collapsed across both groups, better efficiency was associated with increased activation on the semantic trials, and these patterns of activation were found within core language regions within the left hemisphere.

Finally, we also examined the relations between efficiency and the patterns of activation associated with the main effect of Age Group (older > younger). The overall model was significant, F(1,36) = 5.72, *p* < .005, R^2^ = 0.34, although no individual predictor was. A step-wise regression indicated that there was a significant relationship between behavior and the interaction term that accounted for a similar amount of variance as the overall model, F(1,36) = 16.35, *p* < .0005, R^2^ = 0.31. Examination of the correlations revealed that the interaction between RT/accuracy and activation to the main effect of Age Group was driven by a marginally significant negative correlation for the younger adults, suggesting that increases in activation were beneficial for younger but not older adults (younger: *r* = -0.39, *p =* .09; older: *r* = 0.05, n.s.).

### Comparisons with our previous experiment

As we did with our behavioral results, to further assess the influence of the distractors on behavioral performance, we conducted a comparison between the fMRI results from the present study and those from our earlier experiment (Diaz et al., 2014). As can be seen from Figure 8, the patterns of activation are overwhelmingly similar. Our comparison of primary interest was whether older adults’ patterns of brain activation differed across the studies. For the semantic condition, there were no regions where older adults from the present study (with distractors) elicited more activation than older adults from our earlier study (no distractors). For the phonological condition, older adults from the present study elicited greater activation than older adults from our earlier study in bilateral caudate and small region in right posterior cingulate.

**Figure 6.**
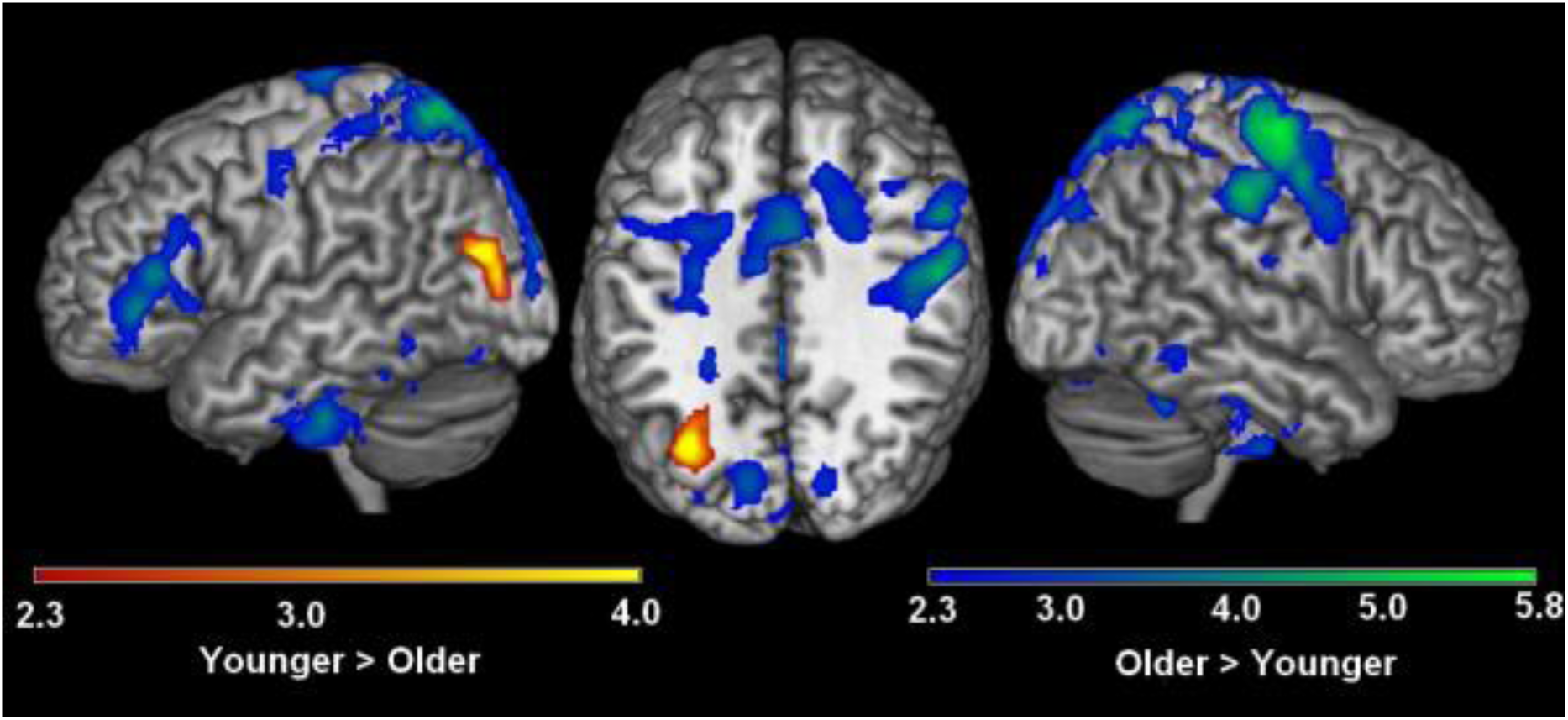
Main Effect of Age. An overview of the regions that comprise the main effect of age are shown. Regions in which younger adults elicited greater activation than older adults are shown in red and regions in which older adults elicited greater activation than younger adults are shown in blue, p < .01 corrected.

**Figure 7.**
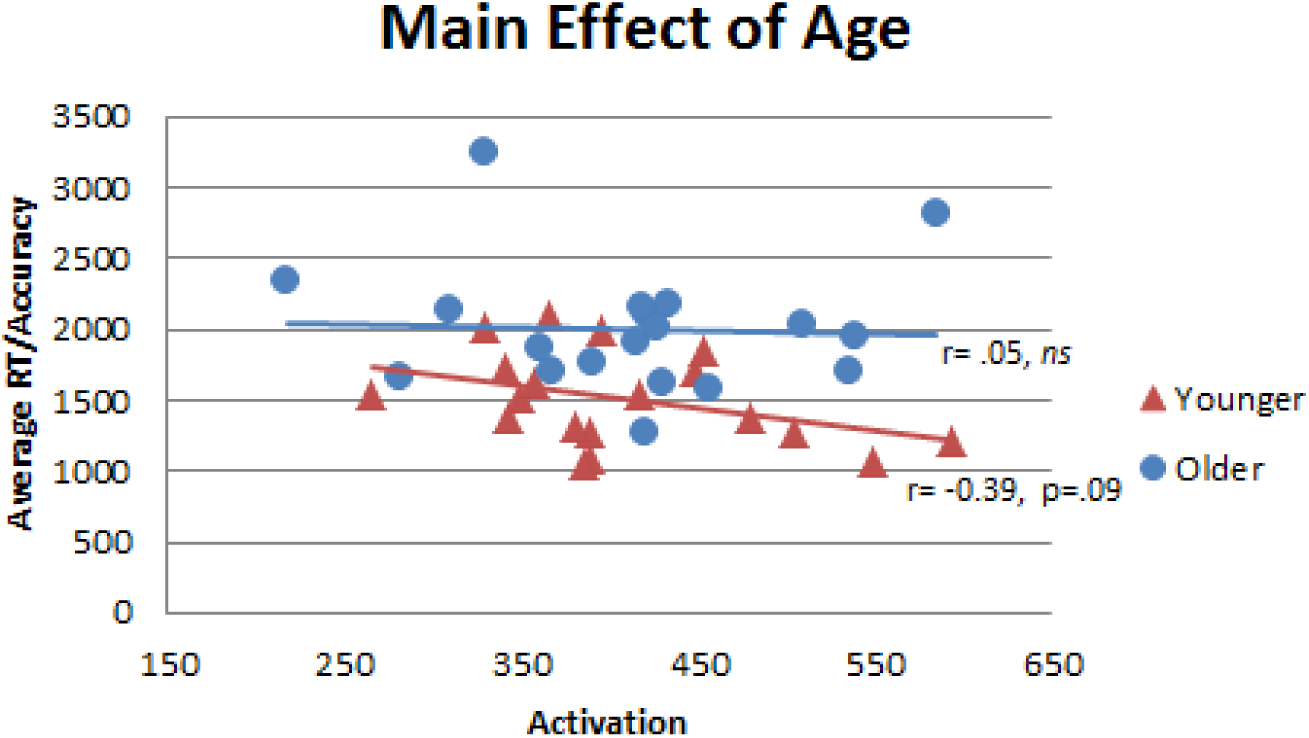
Scatterplot of the Main Effect of Age by group A scatterplot of the relationship between our behavioral measure of RT/accuracy and parameter estimates of the fMRI activation from the main effect of Age are shown. There was a marginally significant negative correlation between behavior and fMRI activation for the younger group only.

**Figure 8.**
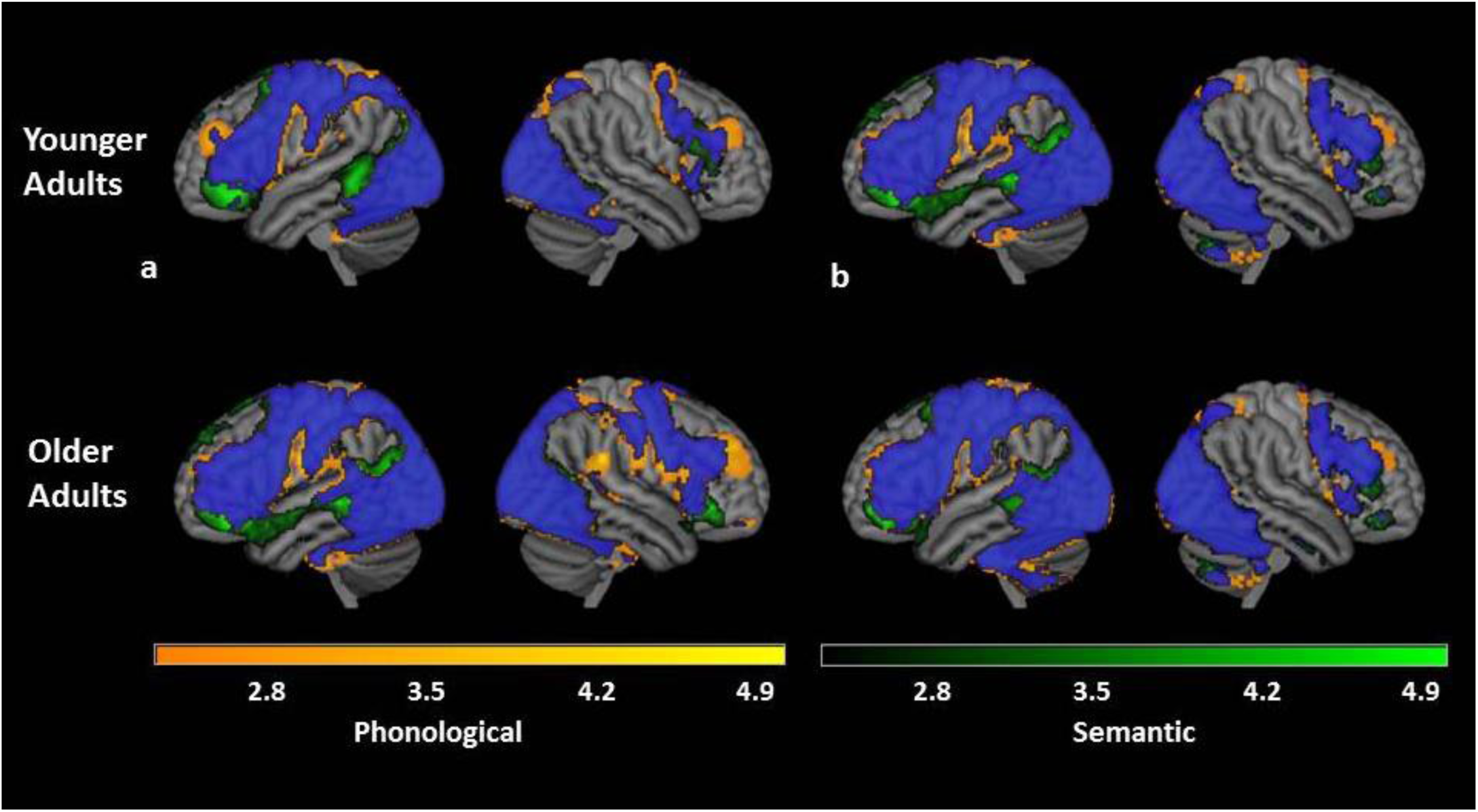
A comparison of results from Diaz et al., 2014 with the current study An overview of the regions that were significantly activated to the phonological (orange) and semantic (green) conditions, are shown broken down by experiment and age group, p < .01 corrected. Results from Diaz et al., 2014 are shown in column a, and results from the present study are shown in column b. Areas of overlap, where both phonological and semantic conditions elicited significant activation are shown in blue.

## DISCUSSION

The present experiment examined the neural and behavioral bases of phonological and semantic processing in older and younger adults. Specifically, we were interested in whether the presence of additional phonological and semantic information would differentially affect processing for younger and older adults. Theoretical perspectives vary on the mechanism behind age-related cognitive declines, with some accounts positing that older adults experience declines in inhibition (Hasher & Zacks, 1988; Lustig, Hasher, & Zacks, 2007) while others suggest that declines are due to age-related differences in transmission of information (Burke et al., 1991). In the present experiment, participants had to decide whether two pictures matched or did not match a phonological, semantic, or perceptual cue in the presence of additional phonologically-, semantically-, or perceptually-related material. Behaviorally, we found a main effect of Age Group in which older adults responded less efficiently than younger adults, and a main effect of Condition, in which phonological trials were responded to less efficiently than semantic trials. In general, older and younger adults responded similarly to semantic and phonological trials, with some age-related slowing on phonological non-match trials. Overall, these results suggest that the presence of additional material did not differentially impair older adults’ performance. While we did observe slower responses from older adults for the phonological non-match trials (i.e., an interaction of Age Group x Condition x Match) which may suggest a differential effect of additional information for older adults (i.e., inhibition deficits), we note that this only occurred for the phonological condition. If older adults were suffering from more general inhibition deficits, we would expect to see age-related differences in performance across all conditions (and perhaps especially for the semantic condition, where well documented cases of semantic inhibition have been reported (e.g., semantic blocking, Belke, Meyer, & Damian, 2005). It is interesting to note that in our first study of semantic and phonological processing in younger and older adults, where we did not include additional semantic or phonological information, we obtained a Condition x Age Group interaction in which older adults were less accurate in processing phonological trials (Diaz et al., 2014). However, in the present study, we found a main effect of Condition in which all participants responded more accurately, if more slowly, to phonological trials. Indeed, explicit comparisons of behavioral performance and patterns of fMRI activation between the two experiments show very few differences. The few differences that were found indicated that older adults from the current study performed more accurately and engaged posterior cingulate to a greater extent for the phonological condition only. These results suggest that the additional phonological information in the present study may have benefitted older adults’ performance.

We also examined brain activation associated with phonological and semantic processing. Consistent with prior research, all language conditions engaged traditional language regions including inferior frontal gyrus, temporal cortex, and parietal cortex, with greater activation in the left hemisphere. Directly comparing semantic and phonological conditions, semantic trials elicited greater activation than phonological trials in well-established regions that have been previously linked to semantic processing including left anterior temporal lobe, middle temporal gyrus, and dorsal medial prefrontal cortex. Phonological trials elicited greater activation than semantic trials in bilateral precuneus and posterior cingulate. While these are not traditional language or phonological regions, the cingulate is involved in task control, and has been linked to language control and monitoring during language production (Abutalebi et al., 2008; Piai, Roelofs, Acheson, & Takashima, 2013). The precuneus has been shown to be involved in imagery and in both auditory and visual retrieval (Fletcher et al., 1995; Huijbers, Pennartz, Rubin, & Daselaar, 2011; Kosslyn, 2003).

In looking at differences across age groups, younger adults elicited greater activation than older adults in left occipital cortex. This specific finding is consistent with what others have reported as a posterior to anterior age-related shift in activation, in which older adults engage posterior regions less than younger adults (Davis, Dennis, Daselaar, Fleck, & Cabeza, 2008). Moreover, older adults elicited greater activation than younger adults in anterior regions including left inferior frontal gyrus, left anterior cingulate cortex, and bilateral pre- and post-central gyrus. However, older adults also elicited greater activation than younger adults in other posterior regions as well including bilateral supramarginal gyrus and precuneus.

What we were most interested in, however, was relating patterns of brain activation to behavioral performance. As noted previously, we did not find large age-related differences in behavior. However, we used regression analyses to examine these brain-behavior relations more thoroughly. Overall, the strongest correlations were found examining patterns of semantic processing. Here we found that across both groups, increases in functional activation in language related regions were associated with increased processing efficiency (i.e., decreases in RT/accuracy scores). Because this effect was found for both groups and in language-related regions, this suggests that older adults semantic processing and engagement of semantic regions is consistent across the lifespan. In contrast, while the overall regression model of age and fMRI activation predicted efficiency in the phonological condition, only younger adults had a marginally significant positive relation between fMRI activation and efficiency. Older adults did not show any significant relation between these variables, and moreover, the additional activation that they produced compared to younger adults was also not related to behavioral performance. Overall, while there was evidence of maintenance of the neural systems supporting semantic processing, the absence of a relationship between brain activation and performance for older adults is consistent with declines in neural efficiency in phonological processing.

While our results shed light on cognitive and neural theories of aging, the present experiment was limited in several ways. We are interested in language production and while our phonological manipulation required covert naming of pictures, there may be some differences between the neural bases of covert and overt production. In particular, our results could not examine age-related differences in articulatory processes. Moreover, while we found main effects of Age Group and Condition, we did not find many interactions with Age Group. It is possible that with a different manipulation stronger age-related differences may emerge.

In conclusion, our results help to shed light on cognitive and neural theories of aging. Inconsistent with the Inhibition Deficit Theory (Hasher & Zacks, 1988), there was no evidence to suggest that older adults were more impaired than younger adults by the presence of additional semantic information or more facilitated by the presence of phonological information. Neurally, we observed consistent engagement of regions during the semantic condition including anterior temporal lobe and dorsal medial prefrontal cortex across the lifespan. Moreover, increases in semantic fMRI activation were associated with increases in response efficiency. However, older adults also exhibited increases in activation in other brain regions that were unrelated to behavioral performance, suggesting an age-related decline in neural efficiency of other brain systems, such as the phonological system, consistent with the Transmission Deficit Hypothesis (Burke et al., 1991).

## Acknowledgments

This project was funded by NIA grant R01 AG034138 (MTD) and R01 AG039684 (DJM). We thank Sarah Danehower for assistance with data analysis and Anna Eppes for assistance with figure preparation. We also thank the staff and scientists at the Duke University Brain Imaging and Analysis Center, where the data were collected, especially the center director Allen W. Song, for their support of this project.

## Disclosure Statement

The authors declare no conflicts of interest.

## Footnotes

1 Although the phonological question involved a letter cue, correct responses required phonological retrieval of the object name because orthographic retrieval is mediated by phonology (e.g., Bonin, Peereman, & Fayol, 2001).

## References

Abutalebi, J., Annoni, J. M., Zimine, I., Pegna, A. J., Seghier, M. L., Lee-Jahnke, H., … Khateb, A. (2008). Language control and lexical competition in bilinguals: an event-related FMRI study. Cerebral Cortex, 18(7), 1496–1505. doi:10.1093/cercor/bhm182

Balota, D. A., Yap, M. J., Cortese, M. J., Hutchison, K. I., Kessler, B., Loftis, B., … Treiman, R. (2007). The English Lexicon Project. Behavior Research Methods, 39, 445–459.

Beckmann, C. F., Jenkinson, M., & Smith, S. M. (2003). General multilevel linear modeling for group analysis in FMRI. Neuroimage, 20(2), 1052–1063.

Belke, E., Meyer, A. S., & Damian, M. F. (2005). Refractory effects in picture naming as assessed in a semantic blocking paradigm. Quarterly Journal of Experimental Psychology, Section A, Human Experimental Psychology, 58(667-692). doi:10.1080/02724980443000142

Bonin, P., Peereman, R., & Fayol, M. (2001). Do phonological codes constrain the selection of orthographic codes in written picture naming? Journal of Memory & Language, 45, 688– 720.

Bowles, N. L., Williams, D., & Poon, L. W. (1983). On the use of word association norms in aging research. Experimental Aging Research, 9, 175–177.

Brown, R., & McNeill, D. (1966). The “tip of the tongue ” phenomenon. Journal of Verbal Learning and Verbal Behavior, 5(4), 325–337.

Burke, D. M., Mackay, D. G., Worthley, J. S., & Wade, E. (1991). On the tip of the tongue: What causes word finding failures in young and older adults? Journal of Memory & Language, 30, 542–579.

Burke, D. M., & Shafto, M. A. (2008). Language and aging. In F. I. M. Craik & T. A. Salthouse (Eds.), The Handbook of Aging and Cognition(3rd ed., pp. 373-443). New York: Psychology Press.

Burke, D. M., White, H., & Diaz, D. L. (1987). Semantic priming in young and older adults: Evidence for age constancy in automatic and attentional processes. Journal of Experimental Psychology: Human Perception and Performance, 13, 79–88.

Cabeza, R., Anderson, N. D., Locantore, J. K., & McIntosh, A. R. (2002). Aging gracefully: Compensatory brain activity in high performing older adults. Neuroimage, 17(3), 1394– 1402. doi:10.1006/nimg.2002.1280

Carlson, M. C., Hasher, L., Zacks, R. T., & Connelly, S. L. (1995). Aging, distraction, and the benefits of predictable location. Psychology & Aging, 10, 427–436.

Christensen, K. J., Moye, J., Armson, R. R., & Kern, T. M. (1992). Health screening and random recruitment for cognitive aging research. Psychology & Aging, 7, 204 – 208.

Connelly, S. L., Hasher, L., & Zacks, R. T. (1991). Age and reading: The impact of distraction. Psychology & Aging, 6, 533–541.

Dale, A. M. (1999). Optimal experimental design for event-related fMRI. Human Brain Mapping, 8, 109–114.

Davis, S. W., Dennis, N. A., Daselaar, S. M., Fleck, M. S., & Cabeza, R. (2008). Que PASA? The posterior-anterior shift in aging. Cerebral Cortex, 18(5), 1201–1209.

Desikan, R. S., Segonne, F., Fischl, B., Quinn, B. T., Dickerson, B. C., Blacker, D., … Killiany, R. J. (2006). An automated labeling system for subdividing the human cerebral cortex on MRI scans into gyral based regions of interest. Neuroimage, 31, 968–980.

Diaz, M. T., Johnson, M. A., Burke, D. M., & Madden, D. J. (2014). Age-related differences in the neural bases of phonological and semantic processes. Journal of Cognitive Neuroscience, 26(12), 2798–2811. doi:10.1162/jocn_a_00665

Fletcher, P. C., Frith, C. D., Baker, S. C., Shallice, T., Frackowiak, R. S., & Dolan, R. J. (1995). The mind’s eye--precuneus activation in memory-related imagery. Neuroimage, 2(3), 195–200.

Friedman, L., & Glover, G. H. (2006). Report on a multicenter fMRI quality assurance protocol. Journal of Magnetic Resonance Imaging, 23(6), 827–839.

Geva, S., Jones, P. S., Crinion, J. T., Price, C. J., Baron, J. C., & Warburton, E. A. (2012). The effect of aging on the neural correlates of phonological word retrieval. Journal of Cognitive Neuroscience, 24(11), 2135–2146. doi:10.1162/jocn_a_00278

Glover, G. H., Mueller, B., Van Erp, T., Liu, T. T., Greve, D., Voyvodic, J., … FBIRN. (2012). Function biomedical informatics research network recommendations for prospective multi-center functional neuroimaging studies. Journal of Magnetic Resonance Imaging, 36(1), 39–54.

Greve, D. N., & Fischl, B. (2009). Accurate and robust brain image alignment using boundary-based registration. Neuroimage, 48(1), 63–72.

Hasher, L., & Zacks, R. T. (1988). Working memory, comprehension, and aging: A review and a new view. In G. H. Bower (Ed.), The Psychology of Learning and Motivation (Vol. 22, pp. 193-225). San Diego, CA: Academic Press.

Hayasaka, S., & Nichols, T. E. (2003). Validating cluster size inference: random field and permutation methods. Neuroimage, 20, 2343–2356.

Horowitz, T. S., & Wolfe, J. M. (2003). Memory for rejected distractors in visual search? Visual Cognition, 10(3), 257–298.

Huijbers, W., Pennartz, C. M., Rubin, D. C., & Daselaar, S. M. (2011). Imagery and retrieval of auditory and visual information: neural correlates of successful and unsuccessful performance. Neuropsychologia, 49(7), 1730–1740.

Jenkinson, M., Bannister, P. R., Brady, J. M., & Smith, S. M. (2002). Improved optimisation for the robust and accurate linear registration and motion correction of brain images. Neuroimage, 17(2), 825–841.

Jenkinson, M., & Smith, S. M. (2001). A global optimisation method for robust affine registration of brain images. Medical Image Analysis, 5(2), 143–156.

Kemper, S., & Sumner, A. (2001). The structure of verbal abilities in young and older adults. Psychology & Aging, 16(2), 312–322.

Kosslyn, S. M. (2003). Understanding the mind’s eye… And nose. Nature Neuroscience, 6(11), 1124–1125. doi:10.1038/nn1103-1124

Kriegeskorte, N., Simmons, W. K., Bellgowan, P. S., & Baker, C. I. (2009). Circular analysis in systems neuroscience: the dangers of double dipping. Nature Neuroscience, 12(5), 535– 540.

Li, S. C., Lindenberger, U., & Sikstrom, S. (2001). Aging cognition: from neuromodulation to representation. Trends in Cognitive Sciences, 5(11), 479–486.

Lustig, C., Hasher, L., & Zacks, R. T. (2007). Inhibitory deficit theory: Recent developments in a new view. In D. S. Gorfein & C. M. MacLeod (Eds.), The Place of Inhibition in Cognition. Washington, D.C.: American Psychological Association.

Madden, D. J., Pierce, T. W., & Allen, P. A. (1993). Age-related slowing and the time course of semantic priming in visual word identification. Psychology & Aging, 8(4), 490–507. doi:10.1037/0882-7974.8.4.490

Oldfield, R. C., & Wingfield, A. (1965). Response latencies in naming objects. Quarterly Journal of Experimental Psychology, 17, 273–281.

Piai, V., Roelofs, A., Acheson, D. J., & Takashima, A. (2013). Attention for speaking: domain-general control from the anterior cingulate cortex in spoken word production. Frontiers in Human Neuroscience, 7, 832. doi:10.3389/fnhum.2013.00832

Shafto, M. A., Stamatakis, E. A., Tam, P. P., & Tyler, L. K. (2010). Word retrieval failures in old age: The relationship between structure and function. Journal of Cognitive Neuroscience, 22(7), 1530–1540. doi:10.1162/jocn.2009.21321

Singer, T., Verhaeghen, P., Ghisletta, P., Lindenberger, U., & Baltes, P. B. (2003). The fate of cognition in very old age: six-year longitudinal findings in the Berlin Aging Study (BASE). Psychology & Aging, 18(2), 318–331.

Smith, S. M. (2002). Fast robust automated brain extraction. Human Brain Mapping, 3, 143-155.

Smith, S. M., Jenkinson, M., Woolrich, M. W., Beckman, C. F., Behrens, T. E. J., Johansen-Berg, H., … Matthews, P. M. (2004). Advances in functional and structural MR image analysis and implementation as FSL. Neuroimage, 23, 208–219.

Townsend, J. T., & Ashby, F. G. (1983). The stochastic modeling of elementary psychological processes. Cambridge, UK: Cambridge University Press.

Verhaeghen, P. (2003). Aging and vocabulary score: A meta-analysis. Psychology & Aging, 18(2), 332–339.

Voyvodic, J. T. (1999). Real-time fMRI paradigm control, physiology, and behavior combined with near real-time statistical analysis. Neuroimage, 10(2), 91–106. doi:10.1006/nimg.1999.0457

Wierenga, C. E., Benjamin, M., Gopinath, K., Perlstein, W. M., Leonard, C. M., Gonzalez Rothi, L. J., … Crosson, B. (2008). Age-related changes in word retrieval: Role of bilateral frontal and subcortical networks. Neurobiology of Aging, 29(3), 436–451. doi:10.1016/j.neurobiolaging.2006.10.024

Woolrich, M. W., Behrens, T. E. J., Beckman, C. F., Jenkinson, M., & Smith, S. M. (2004). Multi-level linear modelling for FMRI group analysis using Bayesian inference. Neuroimage, 21(4), 1732–1747.

Woolrich, M. W., Jbabdi, S., Patenaude, B., Chappell, M., Makni, S., Behrens, T. E. J., … Smith, S. M. (2009). Bayesian analysis of neuroimaging data in FSL. Neuroimage, 45, S173–186.

Woolrich, M. W., Ripley, B. D., Brady, J. M., & Smith, S. M. (2001). Temporal autocorrelation in univariate linear modelling of FMRI data. Neuroimage, 6, 1370–1386.

